# A super sensitive auxin-inducible degron system with an engineered auxin-TIR1 pair

**DOI:** 10.1101/2020.01.20.912113

**Authors:** Kohei Nishimura, Ryotaro Yamada, Shinya Hagihara, Rie Iwasaki, Naoyuki Uchida, Takumi Kamura, Koji Takahashi, Keiko U. Torii, Tatsuo Fukagawa

## Abstract

<u>A</u>uxin-<u>I</u>nducible <u>D</u>egron (AID) technology enables conditional depletion of targeted proteins. However, the applicability of the AID in vertebrate cells has been limited due to cytotoxicity caused by high auxin concentrations. Here, we establish an improved AID system using an engineered orthogonal auxin-TIR1 pair, which exhibits over 1,000 times stronger binding. With ~1,000-fold less auxin concentration, we achieved to generate the AID-based knockout cells in various human and mouse cell lines in a single transfection.

## Main

Gene knockout is a critical method to examine the functions of gene products. For essential genes, however, conditional knockout methods must be used due to cell lethality. Transcription of a target gene can be turned off conditionally under control of a conditional promoter such as a tetracycline responsive promoter. However, it takes long time (usually more than two days) to deplete a target product after turning off transcription. By contrast, auxin-inducible degron (AID) technology enables direct and conditional depletion of a target gene product. A phytohormone auxin (Indole-3-acetic acid, IAA) is used as a degradation inducer in the AID. In non-plant cells expressing an auxin receptor TRANSPORT INHIBITOR RESPONSE1 (TIR1), TIR1 is recruited to Skp1, Cul1 and Rbx1 to form a chimeric E3 ubiquitin ligase, SCF^TIR1^ complex. A target protein fused with IAA17 (called AID-tag) is degraded in an auxin dependent manner via ubiquitin proteasome pathway in non-plant cells expressing TIR1^1^. The AID technology has been widely used to generate conditional knockout yeasts and vertebrate cell lines ^1^ ^2^. For example, this technique was recently applied to two essential genes, encoding condensin I and II, which play distinct roles in mitotic chromosome organization ^3^.

To generate AID-based conditional knockout cell lines, a target gene fused to the AID-tag sequence must be integrated into the endogenous target gene loci by homologous recombination, in cells expressing TIR1 ^4^. However, in some cancer cell lines, such as HeLa cells, which have multiple sets of chromosomes, it is difficult to edit all the endogenous target alleles. To overcome this disadvantage, we previously developed an efficient method to generate AID-based conditional knockouts in chicken DT40 cells. In that method, CRISPR/Cas9-based gene targeting ^5^ ^6^ was used, combined with integration of an AID plasmid (pAID) possessing the rice (*Oryza sativa*) TIR1 gene (*OsTIR1*) and the target protein with an AID-tag (Fig. 1a) ^7^. This method is simple and can be performed by a single transfection to generated AID-based conditional knockout cells (see below). However, the application of this method to other cell lines has not been tested yet.

**Fig. 1.**
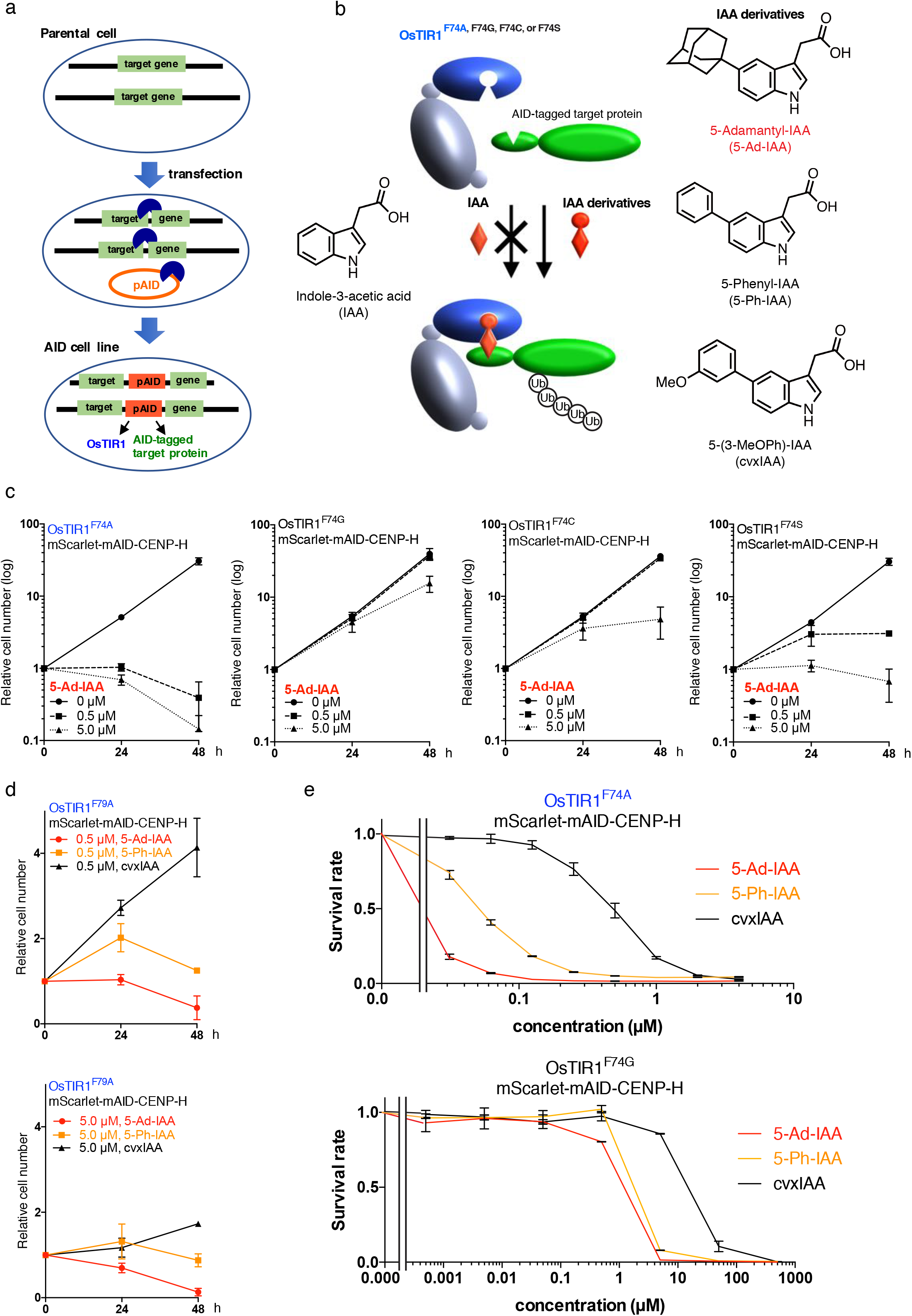
Optimization of an improved AID system with OsTIR1 F74 mutant and IAA derivatives in chicken DT40 cells. **a**, Schematic illustration of a single-step method to generate an AID-based knockout cell line, in which CRISPR/Cas9-based gene targeting is coupled with AID-plasmid integration to express both OsTIR1 and an AID-tagged target protein. Parental cells are transfected simultaneously with three different plasmids: (i) The pAID plasmid encoding OsTIR1, an AID-tagged target protein, and a protein that confers resistance to the drug blasticidin; (ii) the pX330 Crispr/Cas9 plasmid for disrupting a target gene; and (iii) the pX330 Crispr/Cas9 plasmid for linearizing the pAID plasmid (see Supplementary Fig. 3 for plasmid constructions). **b**, Schematic illustration of a “bump and hole” AID system with a high-affinity combination of an OsTIR1F74 mutant and IAA derivatives. The OsTIR1F74 mutant, a functional subunit of the Skp, Cullin, F-box containing (SCF) E3 ubiquitin ligase complex, efficiently interacts with an AID-tagged target protein through IAA derivatives, including 5-Ad-IAA but not conventional IAA, to ubiquitinate the target protein for rapid proteasome-dependent degradation. Green shows a target protein fused with an AID-tag. Gray shows subunits of the SCF complex, except for TIR1. **c**, Growth curves of various AID-based CENP-H knockout cell lines expressing either OsTIR1F74A, F74G, F74C, or F74G in the presence of various concentrations of 5-Ad-IAA. **d**, Growth curves of these cells are shown at 0.5 or 5 µM of three different AID derivatives (5-Ad-IAA, 5-Ph-IAA, and cvxIAA). Error bars indicate the standard deviation of three independent experiments. **e**, Survival rates of the AID cells with OsTIR1F74A or OsTIR1F74G at various concentrations of three AID inducers (5-Ad-IAA, 5-Ph-IAA, or cvxIAA). Error bars indicate the standard deviation of three independent experiments.

More importantly, a major drawback of the AID technology is that protein-degradation does not occur efficiently at 37 °C, which is an optimal temperature for mammalian cell cultures ^1^. This is because the auxin receptor TIR1 is of plant origin, and most plants grow at temperatures lower than 37°C. To compensate for the inefficient degradation at 37°C, a high concentration (500 µM) of IAA is used in the conventional AID technology ^1^. However, we found that IAA at 500 µM is cytotoxic toward some human cell lines (Supplementary Fig. 1).

To overcome auxin cytotoxicity, we took advantage of an orthogonal auxin-TIR1 system engineered via “bump and hole” approach. Here, a high-affinity interaction occurs pairwise between an auxin derivative with an extra bump, 5-adamantyl-IAA (5-Ad-IAA, also known as pico-cvxIAA), and a modified *Arabidopsis* TIR1 protein with a complementary cavity in the auxin binding pocket achieved by an F79A amino acid (aa) substitution (AtTIR1^F79A^) ^8^ ^9^. First, we performed a yeast two-hybrid assay to determine the effective concentrations of 5-Ad-IAA for the interaction of TIR1 with the auxin-responsive protein *Arabidopsis thaliana* IAA17 (AtIAA17) that was used as the AID-tag (Supplementary Fig. 2). For this assay, we used the full-length AtIAA17 or the minimized form of AtIAA17 (aa 65–132, mAID) ^10^ ^11^, and four different TIR1 proteins (AtTIR1^WT^, AtTIR1^F79A^, OsTIR1^WT^, and OsTIR1^F74A^) at 30 or 37°C (Supplementary Fig. 2): i). The interactions were more than 1000 times stronger using the modified TIR1s (AtTIR1^F79A^ or OsTIR1^F74A^) than with the wild-type TIR1s (AtTIR1^WT^ or OsTIR1^WT^); ii) 5-Ad-IAA was 10000 times effective than IAA; and (iii) interactions were detected at 37 °C, although they were more efficient at 30 °C in yeast. Both AtTIR1^F79A^ and OsTIR1^F74A^ interact with AtIAA17 at low concentrations of 5-Ad-IAA (0.1 to 10 nM), which are approximately 50,000 times lower than that used in the conventional AID technology (500 µM IAA).

Next, we used a combination of the modified TIR1 and the auxin derivative in vertebrate cell lines. We used DT40 cells and selected centromere protein H (CENP-H) as the target protein to test the efficiency of protein depletion. CENP-H is essential for cell growth ^12^; therefore, we could easily evaluate CENP-H degradation by monitoring the growth curves and survival rates of the AID-based CENP-H knockout cells. Using the improved method, we generated AID-based CENP-H conditional knockout DT40 cells with various combinations of the modified TIR1s and auxin derivatives (Fig. 1b and Supplementary Fig. 3).

We first compared OsTIR1^F74A^ with AtTIR1s (AtTIR1^F79A^ or AtTIR1^F79A,D170E,M473L^) with various concentrations of 5-Ad-IAA (Supplementary Fig. 4). D170E and M473L mutations were recently suggested to enhance the affinity of AtTIR1 for the *Arabidopsis* IAA7 protein via auxin ^13^; therefore, we further modified AtTIR1^F79A^ with D170E and M473L mutations and used the full-length IAA17 as an AID-tag. The growth rates of the tested lines, as well as the protein degradation profile based on immunoblots, indicate that OsTIR1^F74A^ degrade CENP-H-IAA17 more efficiently than the AtTIR1s in the presence of 5-Ad-IAA (Supplementary Fig. 4a and b). Under these conditions, CENP-H-IAA17 degradation occurred even at 0.005 µM 5-Ad-IAA (Supplementary Fig. 4b). These results strongly suggest that the growth defects of the CENP-H knockout cells correlate well with the CENP-H protein degradation profiles. Therefore, we conclude that we could evaluate CENP-H degradation by monitoring the growth curves of the AID-based CENP-H knockout cells.

We then tested other amino-acid substitutions of F74 in OsTIR1 (F74G, F74C, and F74S), in addition to F74A (Fig. 1c). For this assay, we used mAID-tagged CENP-H (mScarlet-mAID-CENP-H). We observed fewer growth defects in cells expressing OsTIR1^F74G^, OsTIR1^F74C^, or OsTIR1^F74S^ than in cells expressing OsTIR1^F74A^ in the presence of 5-Ad-IAA (Fig. 1c), which was consistent with a previous finding that AtTIR1^F79G^ and AtTIR1^F79S^ show lower affinities to auxin derivatives than AtTIR1^F79A^ ^8^ ^9^. Therefore, OsTIR1^F74A^ most efficiently degrades mScarlet-mAID-CENP-H in the presence of the 5-Ad-IAA in DT40 cells.

We further tested other auxin derivatives 5-(3-MeOPh)-IAA (cvxIAA) and 5-Phenyl-IAA (5-Ph-IAA)). As shown in Fig. 1d and e, 5-Ad-IAA showed the highest growth inhibition of DT40 cells expressing mScarlet-mAID-CENP-H and OsTIR1^F74A^. We also confirmed that the effective concentrations of all the auxin derivatives were 100 times lower for OsTIR1^F74A^ than for OsTIR1^F74G^ (Fig. 1e). Taken together, we conclude that the combination of OsTIR1^F74A^ and 5-Ad-IAA is the most efficient at degrading CENP-H and that 0.5–5 µM 5-Ad-IAA is sufficient for CENP-H degradation in AID-based CENP-H knockout DT40 cells.

Next, we expanded our improved AID system for other target proteins. For this purpose, we choose other centromeric proteins because of their highly stable property, which make it difficult to deplete them. We generated the improved AID-based knockout DT40 cell lines for CENP-A or CENP-T (Fig. 2a and Supplementary Fig. 5). Based on immunoblots, CENP-A, CENP-T, and CENP-H proteins were undetectable within 2 h after the addition of 5 µM 5-Ad-IAA, but not after the addition of IAA (Fig. 2a and Supplementary Fig. 5a). Efficient degradation of the target proteins was also confirmed by fluorescent microscopy, in which mScarlet fluorescence signals of the target proteins (CENP-A, CENP-H, and CENP-T) became undetectable at 3 h after adding 5 µM 5-Ad-IAA. In contrast, fluorescent signals remained detectable after adding 5 µM IAA (Fig. 2b).

**Fig. 2.**
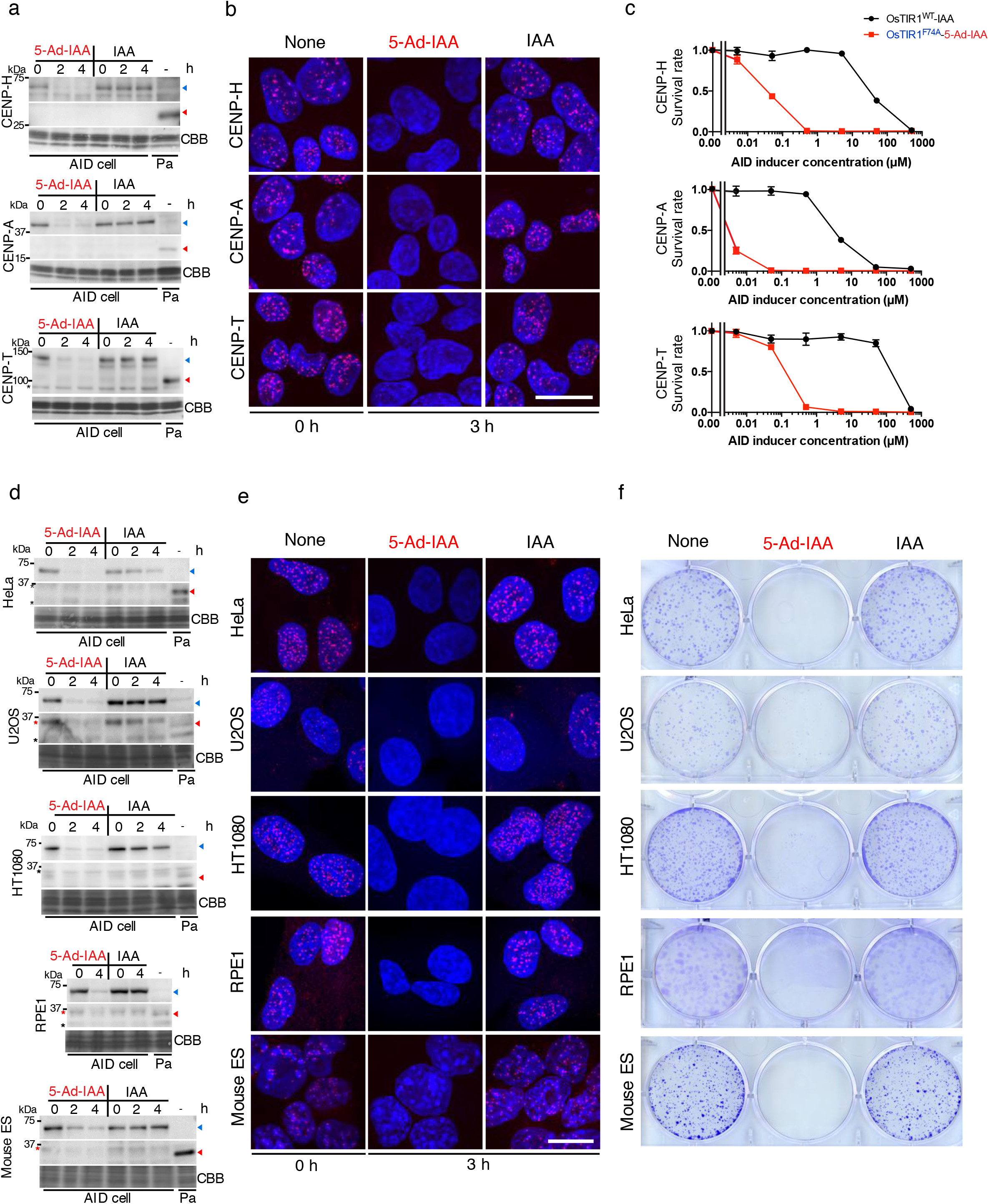
Versatility of the improved AID system. **a**, Immunoblots for the target CENP proteins (CENP-H, CENP-A, and CENP-T) to evaluate their degradation levels in the OsTIR1^F74A^-AID-based knockout cells at 0, 2, and 4 h after adding 5 µM 5-Ad-IAA or 5 µM IAA. Pa, parental cells; blue arrowheads, mScarlet-mAID-CENPs; red arrowheads, endogenous CENP proteins; and asterisks, non-specific signals. The uncropped gel images are shown in Supplementary Fig. 5a. **b**, Fluorescent images of cells expressing the mScarlet-mAID-tagged CENP proteins (CENP-H, CENP-A, and CENP-T) and OsTIR1^F74A^ to evaluate their degradation at 3 h after adding 5 µM 5-Ad-IAA, 5 µM IAA, or the solvent dimethyl sulfoxide (DMSO) (none). Red shows Scarlet signals and blue shows 2-(4-amidinophenyl)-1H-indole-6-carboxamidine (DAPI) signals. Scale bar, 20 μm. **c**, Comparison of the AID-system using OsTIR1^F74A^ and 5-Ad-IAA with the conventional AID system using OsTIR1 and IAA. CENP-H, CENP-A, and CENP-T knockout cell lines were used for this comparison. The relative viabilities of these cells are shown at various concentrations of IAA or 5-Ad-IAA. Error bars indicate the standard deviation of three independent experiments. **d**, Immunoblots for the target protein CENP-H in various OsTIR1^F74A^-AID-based knockout cells after adding 5 µM 5-Ad-IAA or 5 µM IAA (see Fig. 2a for details). The uncropped gel images are shown in Supplementary Fig. 5b. **e**, Fluorescent images of cells expressing mScarlet-mAID-tagged CENP-H and OsTIR1^F74A^ after adding 5 µM 5-Ad-IAA, 5 µM IAA, or the solvent DMSO. Red shows Scarlet signals and blue shows DAPI signals. Scale bar, 20 μm. **f**, Colony-formation assay of the OsTIR1^F74A^-AID-based knockout cells that were cultured for 1 to 2 weeks in the presence of 5 µM 5-Ad-IAA, 5 µM IAA, or the solvent DMSO. Colonies were stained with crystal violet.

The survival rates of the CENP-A, CENP-H, and CENP-T knockout cells varied; however, all cells died at 48 h after the addition of at least 0.5 µM 5-Ad-IAA (Fig. 2c). On the other hand, in cells expressing conventional OsTIR1, the IAA concentrations required for complete cell death were 500 µM for CENP-H or CENP-T knockout cells and 50 µM for CENP-A knockout cells. These results emphasize that the improved AID system works for other target proteins with extremely low concentrations of 5-Ad-IAA in chicken DT40 cells.

Finally, we applied the improved AID system to other cell lines, including human cultured cell lines (HeLa, HT1080, U2OS, and RPE1) and mouse embryonic stem (ES) cells. Using mScarlet-mAID-CENP-H as the target protein, we generated AID-based conditional knockout cell lines expressing OsTIR1^F74A^ in the presence of 5 µM 5-Ad-IAA (Fig. 2d, e, f and Supplementary Fig. 5b).

In each of the tested lines, immunoblot analyses showed that mScarlet-mAID-CENP-H was degraded in 2–4 h after adding 5 µM 5-Ad-IAA, but not after adding 5 µM IAA (Fig. 2d and Supplementary Fig. 5b). This was further confirmed by microscopy (Fig. 2e), in which mScarlet fluorescence signals of the target protein were not observed in nuclei at 3 h after adding 5 µM 5-Ad-IAA, but were observed at 3 h after adding 5 µM IAA. Consistent with degradation profiles of the essential protein CENP-H, these cell lines showed growth defects and did not form any colonies in the presence of 5 µM 5-Ad-IAA, whereas they formed colonies in the presence of 5 µM IAA (Fig. 2f). These results indicated that AID-based CENP-H conditional knockout lines could be generated from the tested mammalian cell lines.

We also generated AID-based conditional knockout lines for CENP-A, CENP-H, and CENP-C from human HeLa S3 cells, using the improved AID system (Supplementary Fig. 6). Immunoblot analyses and microscopy observations showed that all the target proteins became undetectable 2 hours after application of 5 µM 5-Ad-IAA, but not 5 µM IAA (Supplementary Fig. 6a and b). Consistent with these results, no colonies were formed in any of the knockout lines in the presence of 5-Ad-IAA (Supplementary Fig. 6c), as was the case with the other mammalian cells (Fig. 2d). Taken together, our results demonstrate that the improved AID system is applicable to various target proteins and cell lines.

In this study, we established a super-sensitive AID system that enables over a thousand-fold reduction in auxin concentration and applicable to a wide variety of cell lines, including cancer cell lines with an aneuploid karyotype. Our method is simple, and only one transfection is sufficient to generate conditional knockout lines for various target proteins. Protein degradation occurs at extremely low concentrations when using 5-Ad-IAA (5 µM) compared with the conventional AID method, which requires high concentration of IAA (500 µM). The improved AID system will be useful to reveal the molecular mechanisms underlying different biological processes in a variety of cell lines.

## Methods

### Cell culture

HeLa cells, HeLa-S3 cells, HT1080 cells, U2OS cells, RPE1 cells are cultured in DMEM (Nakarai) supplemented with 10% FBS, 100U/ml of penicillin/streptomycin (Nakarai) at 37°C in 5% CO_2_. DT40 cells are cultured in DMEM (Nakarai) supplemented with 10% FBS, 1% chicken Serum(Gibco), 100U/ml of penicillin/streptomycin (Nakarai) and 50µM of 2-Mercaptethanol (Sigma)at 38.5°C in 5% CO_2_. Feeder Free Mouse ES cells (E14tg2a) are cultured on Gelatinized dish in DMEM (Nakarai) supplemented with 10% FBS, 5% Knockout Serum Replacement, L-glutamine (2 mM), penicillin/streptomycin (100U ml^−1^ each), non-essential amino acid and LIF protein at 37°C in 5% CO_2_.

### Plasmid construction

pAID plasmid was generated by assembling PCR fragments by Infusion(Takara) based clonging from pAID (Addgene: no 72835) with codon optimized OsTIR1. Each cDNA was cloned to EcoRV site of pAID plasmid by In fusion(Takara) cloning.

### Transfection and cloning colonies for AID cell line

4µg of Plasmid DNA (pAID plasmid: pX330 for target gene: p330 for pAID linearlization = 1: 2: 1) was diluted with 50µl of 20mM Glutamic acid (pH4.0).

18 µg of Polyethilenimin(PolySciences) was diluted diluted with 50µl of 20mM Glutamic acid (pH4.0). DNA solution was mix with Polyethilenimin solution and voltex for 5 sec. Mixture solution incubate at room temperature for 15 min. 1 × 10^6 cells (HeLa, U2OS, HT1080, E14tg2a) was diluted with 300 µl of serum free DMEM. Cells were mix with DNA-Polyethilenimin solution and voltex for 5sec.

After 15 min incubation at room temperature, cells were plated to 1, 1/5, 1/25 or 1/125 to 10cm plate. Selection was start with 10 µg /ml of Blastsidin at 24 hrs after the transfection. Selection media was changed every 3 days. Surviving colonies were isolated at 1 – 2 weeks after selction start.

25µg of plasmid DNA (pAID plasmid: pX330 for target gene: p330 for pAID linearlization = 1: 2: 1) was peletted by ethanol precipitation. Pelleted DNA was resuspended with 125µl of R buffer of Neon transfection system (thermos fisher).

2 × 10^5 DT40 cells were diluted in DNA solution and Neon transfection was performed at 1400V, 5 msec, 6 times, 2days after transfection, Cells were Plated to 96 well plate and selection was start at 30 µg/ml of Blastsidin. 1 week after selection start, surviving colonies picked up.

25µg of plasmid DNA (pAID plasmid: pX330 for target gene: p330 for pAID linearlization = 1: 2: 1) was peletted by ethanol precipitation. Pelleted DNA was resuspended with 125µl of R buffer of Neon transfection system (thermos fisher).

2 × 10^5 RPE1 cells were diluted in DNA solution and Neon transfection was performed at 1400V, 30msec, 1times, 2days after transfection, cells were plated to 1, 1/5, 1/25 or 1/125 to 10cm plate. Selection was start with 10 µg /ml of Blastsidin at 24 hrs after the transfection. Selection media was changed every 3 days. Surviving colonies were isolated at 1 – 2 weeks after selection start.

### Fluorecent microscopy

Cells were fixed with 3% Paraphormaldehyde. After DAPI staining, cells were washed with PBS twice and mounted with Vectashield mounting reagent (Vector Laboratories). FISH images were captured by sCMOS camera (Zyla 4.2; Andor) mounted on ECLIPSE Ti microscope (Nikon) with an objective lens (Plan Apo lambda 100×/1.45 NA; Nikon) and CSU-W1 confocal scanner unit (Yokogawa) controlled by NIS elements (Nikon). Z stacked images

### Cell growth assay and Cell viability assay

1 × 10^5 cells / ml of DT40 cells were cultured with or without auxin derivative. Cell concentrations are calculated by Countess II (thermos fisher).

Cell viability was calculated with beta-Glo assay (Promega)

### Colony formation assay

500 – 1000 cells were plated to 6 well plated and cultured at cultured conditions for 1 to 2 weeks. Cells were fixed with cold 100% methanol for 2 min and airdry.

Cells were stained Crystal Violette and washed with water.

### Western blotting assay

Cells were lysed in 1 × Laemmli buffer and incubate 95 oC for 5 min. After sonication, Samples were loaded onto 5%-15% or 5% – 20% gradient gel, activated after running and transferred to PVDF membrane. Membranes were blocked 3% BSA at room temperature for 0.5 – 1 hr and incubate with first antibody (Rabbit anti-chiken CENP-A, Rabbit anti-chicken CENP-H, Rabbit anti-chicken CENP-T, mouse anti-human CENP-A, mouse anti-human CENP-H(sc-365222, SANTA CRUZ), guinia pig anti-human CENP-C or mouse anti-mouse CENP-H(sc-136403, SANTA CRUZ) at 4oC for overnight or at room temperature for 1 hour. After 3 times wash with TBST, membranes were incubated with goat anti-Rabbit IgG, goat anti-mouse IgG or Goat anti-guinia pig IgG antibodies. Detections was performed with ECL Prime(GE healthcare)and all images were acquired with a ChemiDoc MP Imaging system.

### Yeast two hybrid assay

Yeast assays were performed according to the previous report (Uchida et al. 2018). Briefly, EGY48 strain was transformed with pSH18-34 (LexA-operon:LacZ reporter), pGLex313-based plasmid (LexA-DNA-binding-domain-fused TIR1 series), and pJG4-5-based plasmid (B42-transcriptional-activator-fused degron series). Transformed strains were grown at 30 °C on agar plates containing minimal SD base (Clontech, cat. 630411) and –His/–Trp/–Ura dropout supplement (Clontech, cat. 630424). Colonies were grown in SD/–His/–Trp/–Ura medium for one night at 30 °C, and then medium was replaced with liquid medium containing minimal SD/Gal/Raf base (Clontech, cat. 630420), –His/–Trp/–Ura dropout supplement, 50 mM Na-phosphate buffer (pH 7.0), 80 mg/ml X-gal (Wako) and compounds. After 3-day incubation at 30 °C or 37 °C, culture were transferred to white 96-well plates and observed.

