## Supplementary Figure 1-6 for "A super sensitive auxin-inducible degron system with an engineered auxin-TIR1 pair"

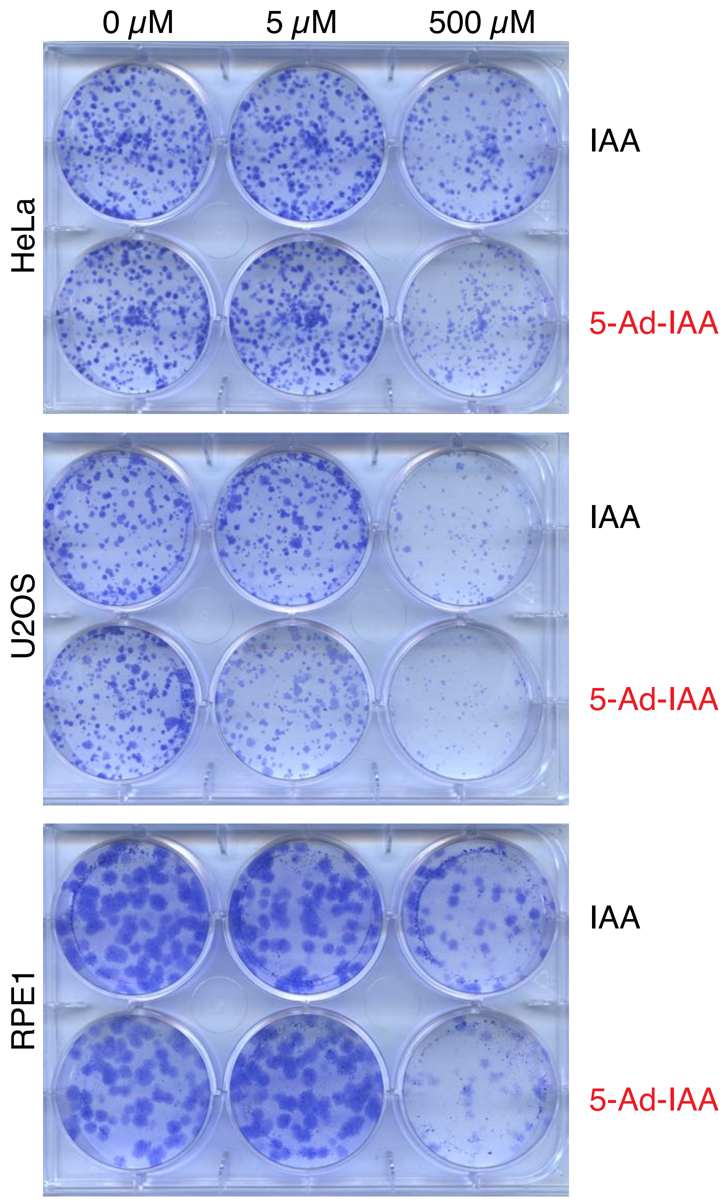


**Supplementary Fig. 1**

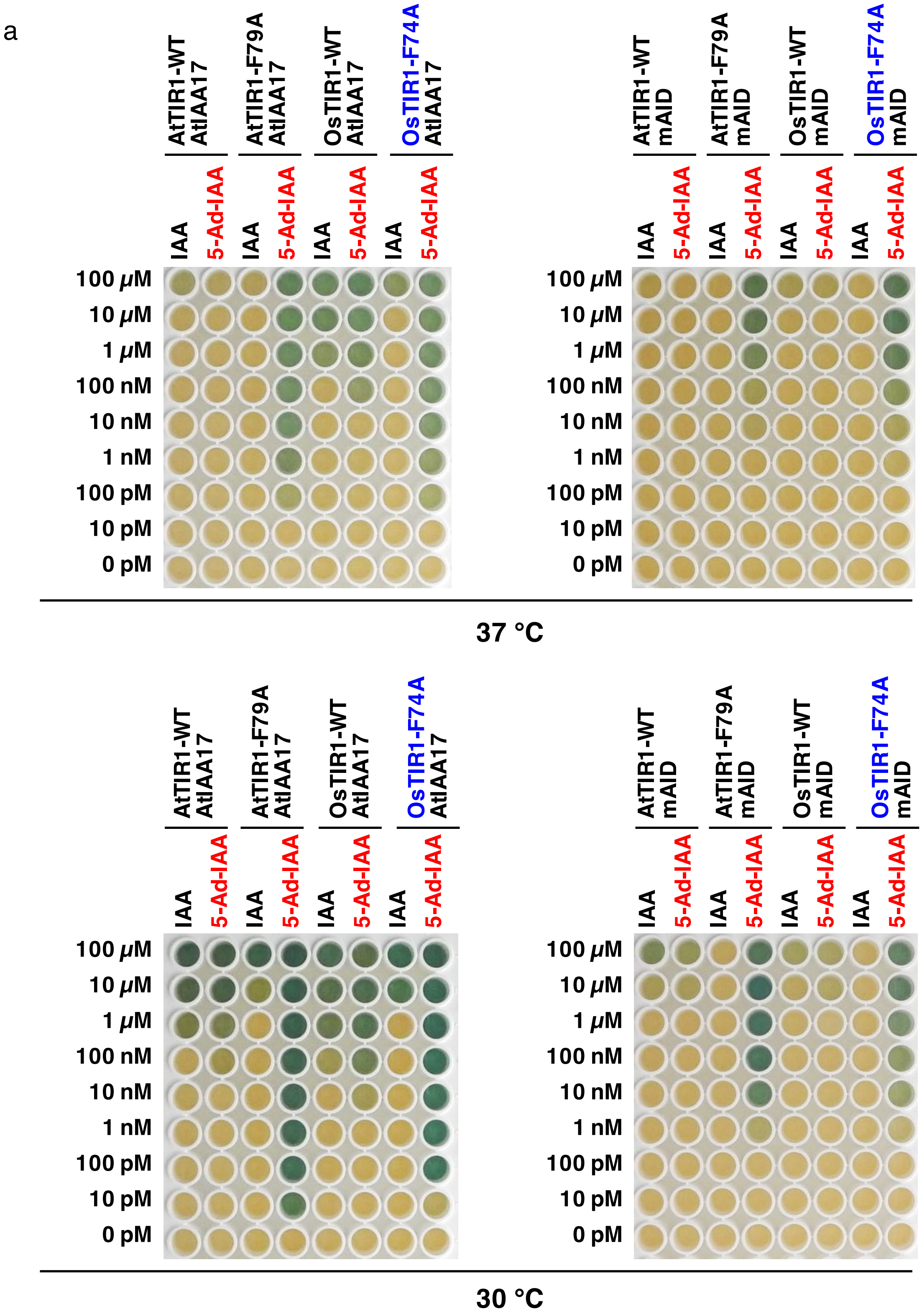


Supplementary Fig. 2

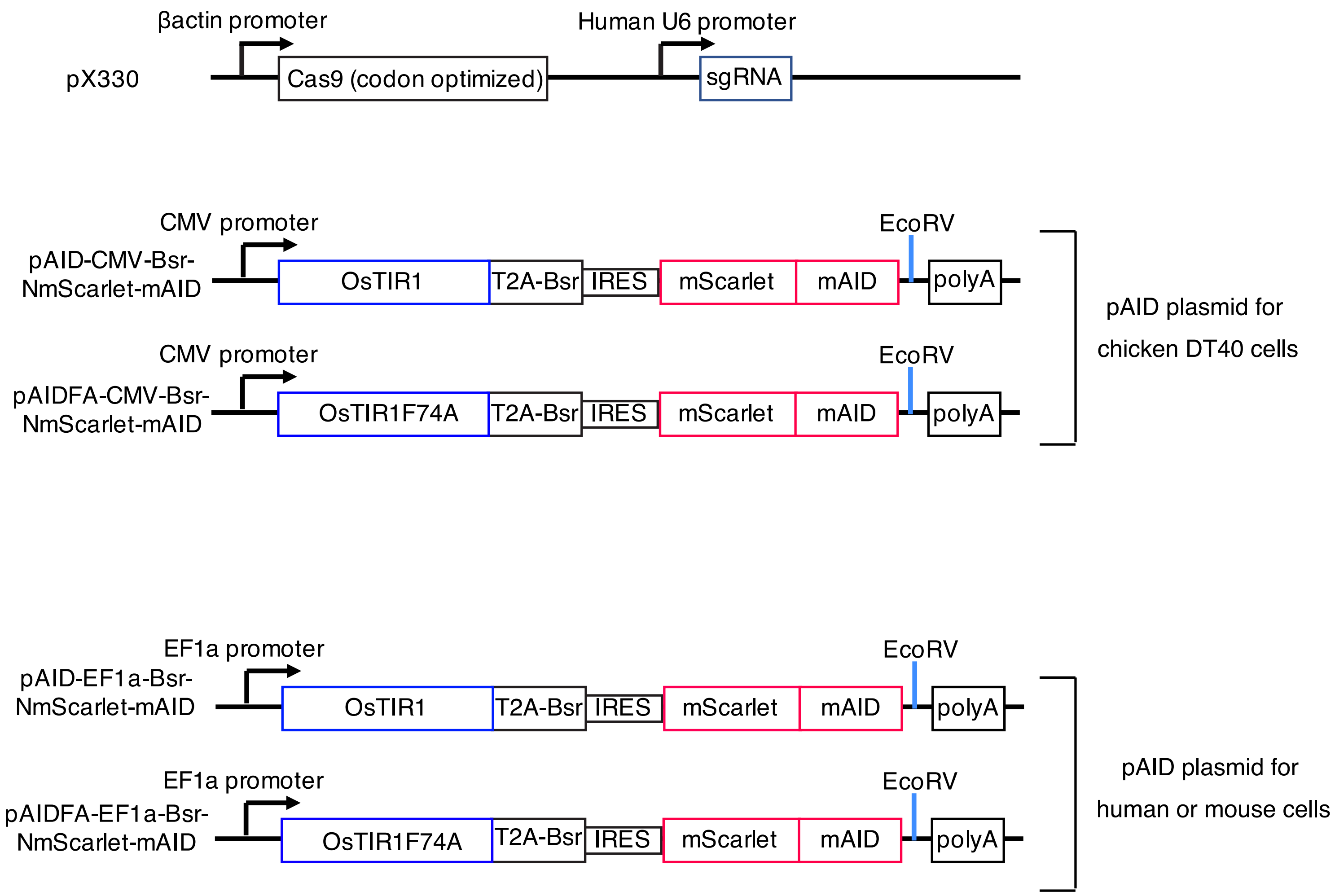


**Supplementary Fig. 3**

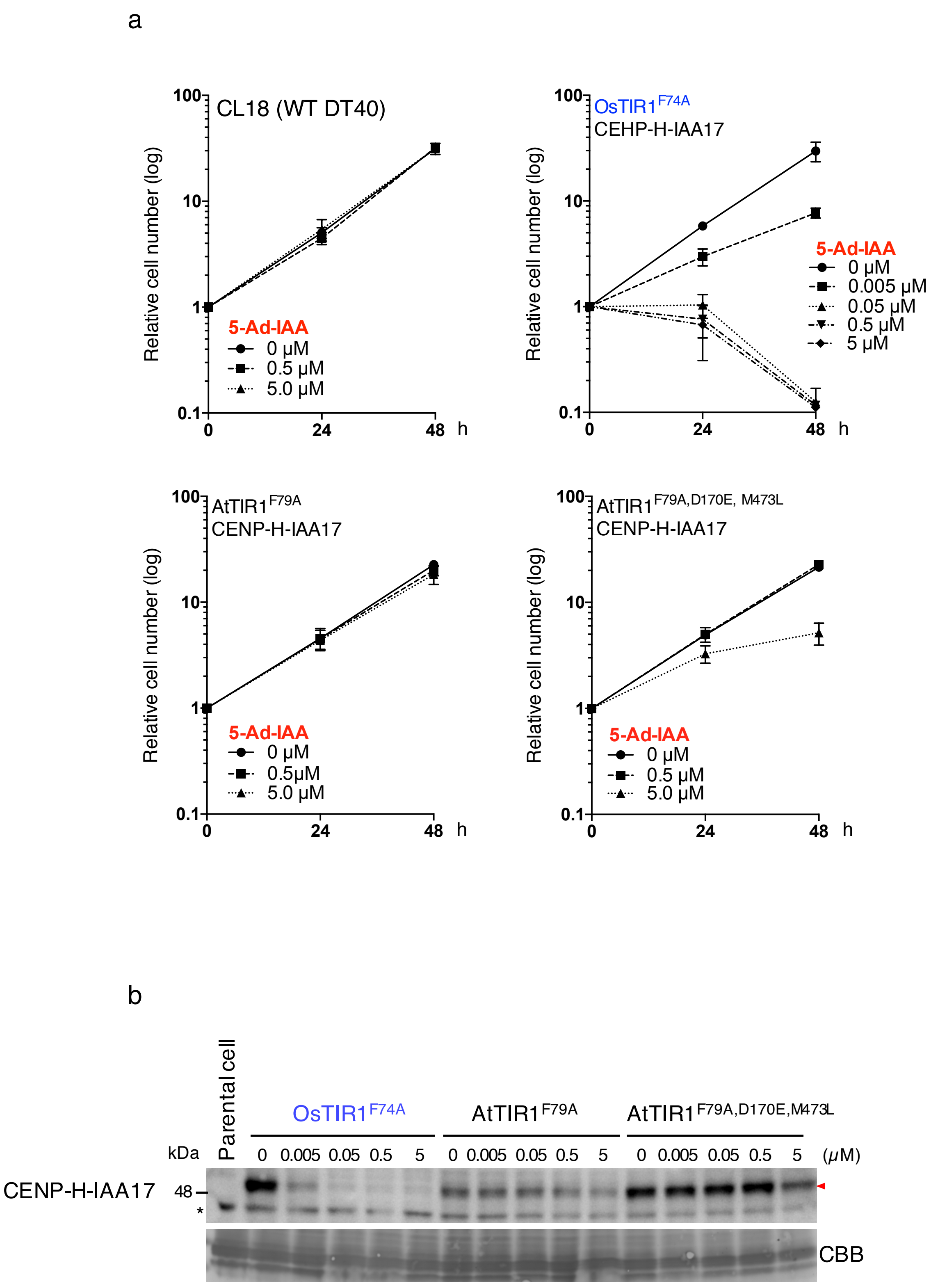


**Supplementary Fig. 4**

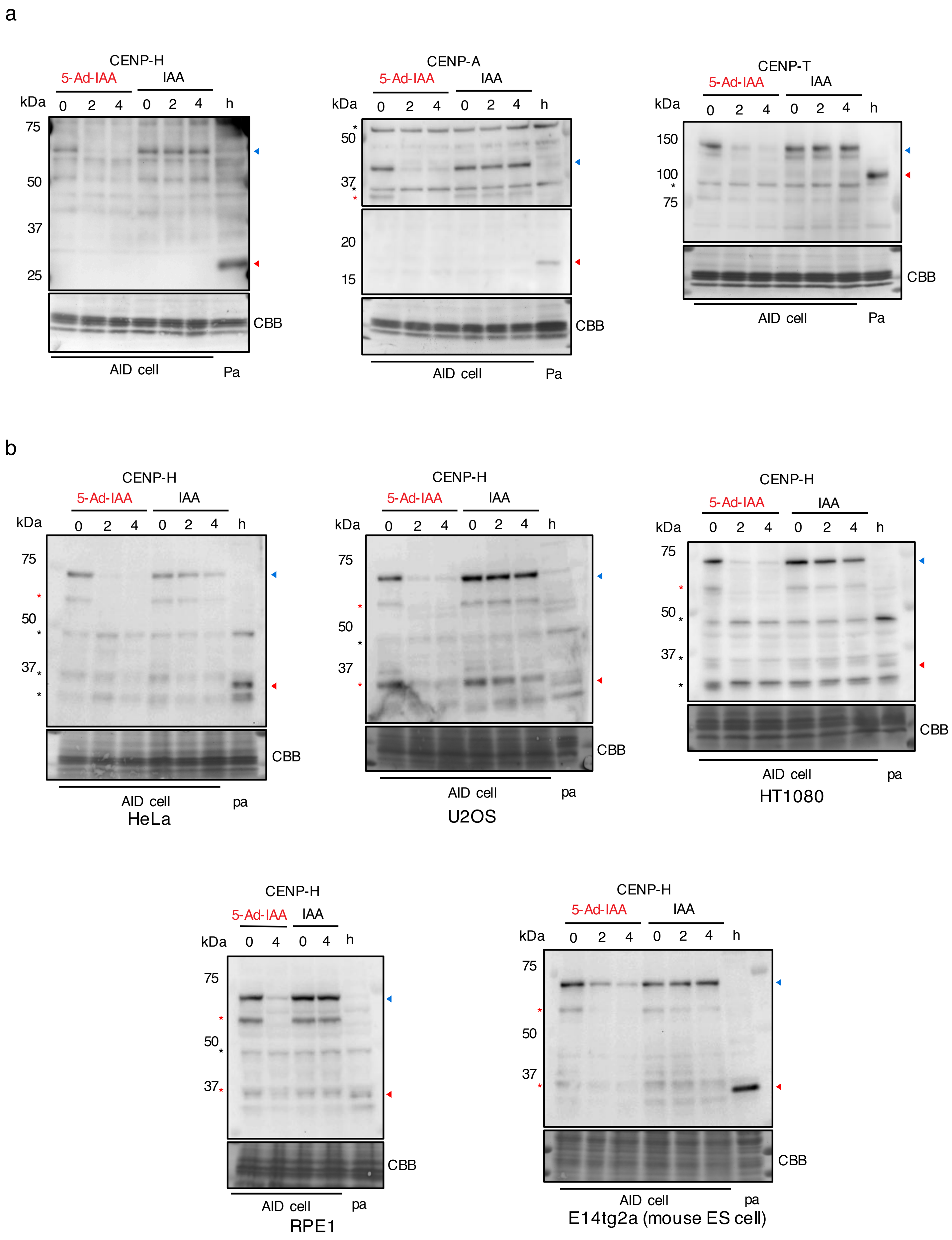


**Supplementary Fig. 5**

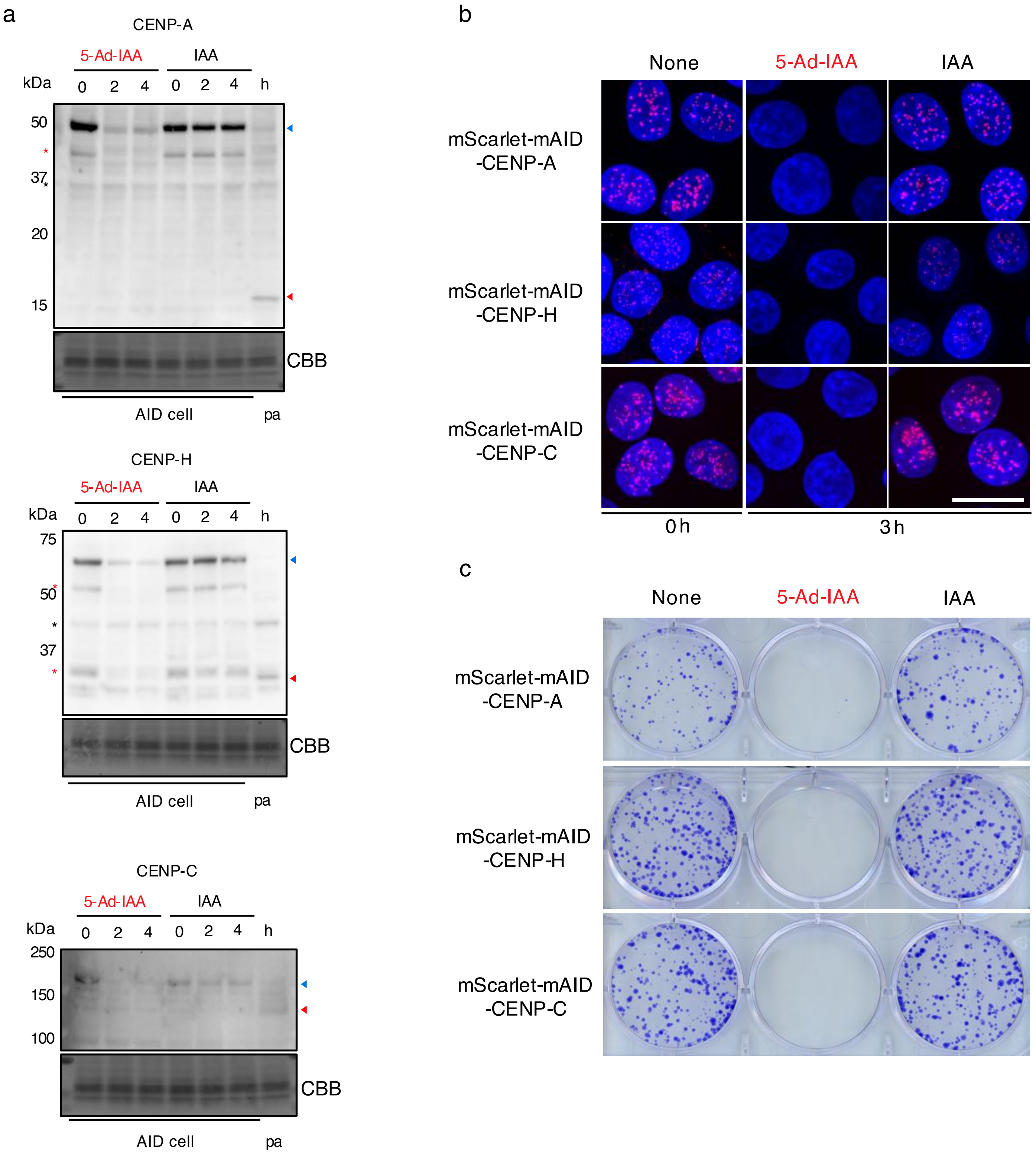


**Supplementary Fig. 6**

**Supplementary Fig. 1. Cytotoxicity of conventional IAA or 5-Ad-IAA.**

Colony-formation assay of HeLa, U2OS and RPE1 cells cultured for 1 to 2 weeks in different concentration of IAA or 5-Ad-IAA. Colonies were stained with crystal violet.

**Supplementary Fig. 2. Yeast-two-hybrid assay showing interaction between TIR1s and the AID-tags to evaluate the optimal concentrations of the AID inducers.**

Four types of TIR1s (AtTIR1$^{WT}$, AtTIR1 $^{F79A}$, OsTIR1$^{WT}$, and OsTIR1$^{F74A}$) were examined for their interactions with the AID-tag (IAA17 or mAID) in the presence of various concentrations of the AID inducers (IAA and 5-Ad-IAA) at 30 or 37 °C.

**Supplementary Fig. 3. Maps of plasmids to generate AID-based knockout cell lines.**

pX330 contains Cas9 and appropriate single guide RNAs (sgRNAs) were cloned. pAID plasmids for DT40 cells contain OsTIR1 and OsTIR1$^{F74A}$ under the control of the Cytomegalovirus (CMV) promoter. cDNA of the target protein was inserted into the EcoRV site using InFusion technology (Takara). pAID plasmids for human or mouse cells contains OsTIR1 and OsTIR1$^{F74A}$ under control of the eukaryotic translation elongation factor 1 Alpha 1 (EF1a) promoter. cDNA of the target protein was inserted into the EcoRV site using the InFusion technology, as was the case for the pAID plasmids in DT40 cells.

**Supplementary Fig. 4. Growth curves and protein levels of the AID-based CENP-H knockout cells expressing either OsTIR1$^{F74A}$, AtTIR1F$^{79A}$, or AtTIR1$^{F79A,D170E,M473L}$**

**a**, Growth curves of parental cells (WT DT40) and the AID-based CENP-H knockout cells expressing either OsTIR1$^{F74A}$, AtTIR1F$^{79A}$ or AtTIR1$^{F79A,D170E,M473L}$ at various concentrations of 5-Ad-IAA. Error bars indicate the standard deviation of three independent experiments.

**b**, Immunoblot for CENP-H-IAA17 in the CENP-H knockout cells expressing either OsTIR1$^{F74A}$, AtTIR1F$^{79A}$, or AtTIR1$^{F79A,D170E,M473L}$ at 3 h after addition of various concentrations of 5-Ad-IAA. The CENP-H-IAA17 position is marked by a red arrowhead and the asterisks show non-specific signals.

**Supplementary Fig. 5. Immunoblot images of target proteins in each AID-based knockout cell.**

**a,** AID-tag fused CENP-H, CENP-A, and CENP-T were detected at 0, 2, and 4 h after adding 5 μM 5-Ad-IAA or 5 μM IAA in each knockout line. Pa, parental cells; blue arrowheads, mScarlet-mAID-CENPs; red arrowheads, endogenous CENP proteins; asterisks, non-specific signals; red asterisks, degradation products. Portions of these immunoblots are shown in Fig. 2a.

**b**, AID-tag fused CENP-H was detected after adding 5 μM 5-Ad-IAA or 5 μM IAA in various mammalian AID-based knockout lines. Portions of these immunoblots are shown in Fig. 2d.

**Supplementary Fig. 6. Application of the improved AID system to HeLa-S3 cells.**

**a**, Immunoblots for the AID-tag fused target proteins (CENP-A, CENP-H, and CENP-C) after adding 5 μM 5-Ad-IAA or 5 μM IAA (see Fig. 1d for details) in each AID-based knockout HeLa S3 cell line expressing OsTIR1$^{F74A}$. Pa, samples from parental cells; blue arrowheads, aid-tagged proteins; red arrowheads, endogenous proteins; asterisks, non-specific signals; red asterisks, degradation products.

**b**, Fluorescent images of AID-based knockout HeLa S3 cells expressing the mScarlet-mAID-tagged CENPs after adding 5 μM 5-Ad-IAA, 5 μM IAA, or the solvent DMSO. Red shows Scarlet signals and blue shows DAPI signals. Scale bar, 20 μm.

**c,** Colony-forming assays of AID-based knockout HeLa S3 cells that were cultured for 1 to 2 weeks in the presence of 5 μM 5-Ad-IAA, 5 μM IAA, or the solvent DMSO. Colonies were stained with crystal violet.
